# Embedding-based alignment: combining protein language models and alignment approaches to detect structural similarities in the twilight-zone

**DOI:** 10.1101/2022.12.13.520313

**Authors:** Lorenzo Pantolini, Gabriel Studer, Joana Pereira, Janani Durairaj, Torsten Schwede

## Abstract

Language models are now routinely used for text classification and generative tasks. Recently, the same architectures were applied to protein sequences, unlocking powerful tools in the bioinformatics field. Protein language models (pLMs) generate high dimensional embeddings on a per-residue level and encode the “semantic meaning” of each individual amino acid in the context of the full protein sequence. Multiple works use these representations as a starting point for downstream learning tasks and, more recently, for identifying distant homologous relationships between proteins. In this work, we introduce a new method that generates embedding-based protein sequence alignments (EBA), and show how these capture structural similarities even in the twilight zone, outperforming both classical sequence-based scores and other approaches based on protein language models. The method shows excellent accuracy despite the absence of training and parameter optimization. We expect that the association of pLMs and alignment methods will soon rise in popularity, helping the detection of relationships between proteins in the twilight-zone.

## 1 Introduction

Protein language models (pLMs) are becoming more popular by the day. These models capture deep “semantic relationships” between different residues in a protein by analyzing their context within the sequence, resulting in neural networks capable of generating meaningful representations at the residue level. These representations, also denoted as embeddings, are vectors in high dimensional space that are used for various downstream machine learning applications; [11] [5]. Recently, pLMs were also leveraged for establishing homologous relationships between sequences. While this is achievable with standard alignment tools [14], whenever the comparison falls into the so-called twilight zone [16], the pairwise signal gets blurry. This is where pLMs shine by capturing relationships way beyond simple sequence comparisons, uncovering otherwise undetected evolutionary relationships that can guide, for example, protein annotation or structure prediction efforts.

Protein sequences are commonly projected into embedding space by averaging the per-residue contributions [8], [7], [17]. However, the meaning of distance in this space is still unclear. In [8] the Euclidean distance in the averaged embedding space was used to quantify sequence similarity, which in turn was used to generate an evolutionary landscape of homologous proteins by connecting sequences to their k-nearest neighbors. The distance between the average representations was used again in [7] to establish distant homology relationships between CATH domains [19]. Performance was improved by contrastive learning to re-project the average embedding representation into a space where similar CATH domains cluster closely together. A similar approach was adopted by [6] to build TM-vec, a tool able to predict TM scores. However, representing a sequence by averaging its per-residue embeddings has limitations. An example is given by multidomain proteins, where a loss of signal can be expected when averaging per-residue embeddings from distinct domains that potentially evolved independently [18]. Furthermore, even for single domain sequences, average-based similarity metrics inherently lose information on order and are affected by comparisons of residues without any evolutionary relationship, which we discuss in section 3.3. An example is shown in figure 1, where the average representation of a sequence is equidistant from a protein with the same structure and another one with a completely different fold. These problems can be approached by methods constructing explicit alignments at the cost of added computational complexity. Two examples of embedding-based alignment methods were introduced by [3] and [6]. In [3], a language model was trained using both sequence and structural information. They showcase a “soft alignment” generated with a weighted sum of all the possible pairwise residue distances. The resulting score predicted homologous relationships between SCOP [2] domains. On the other hand, in [13] a network fed with residue embeddings was trained on protein structures to generate dynamic alignment parameters, such as the score and gap penalty matrices.

**Fig. 1:**
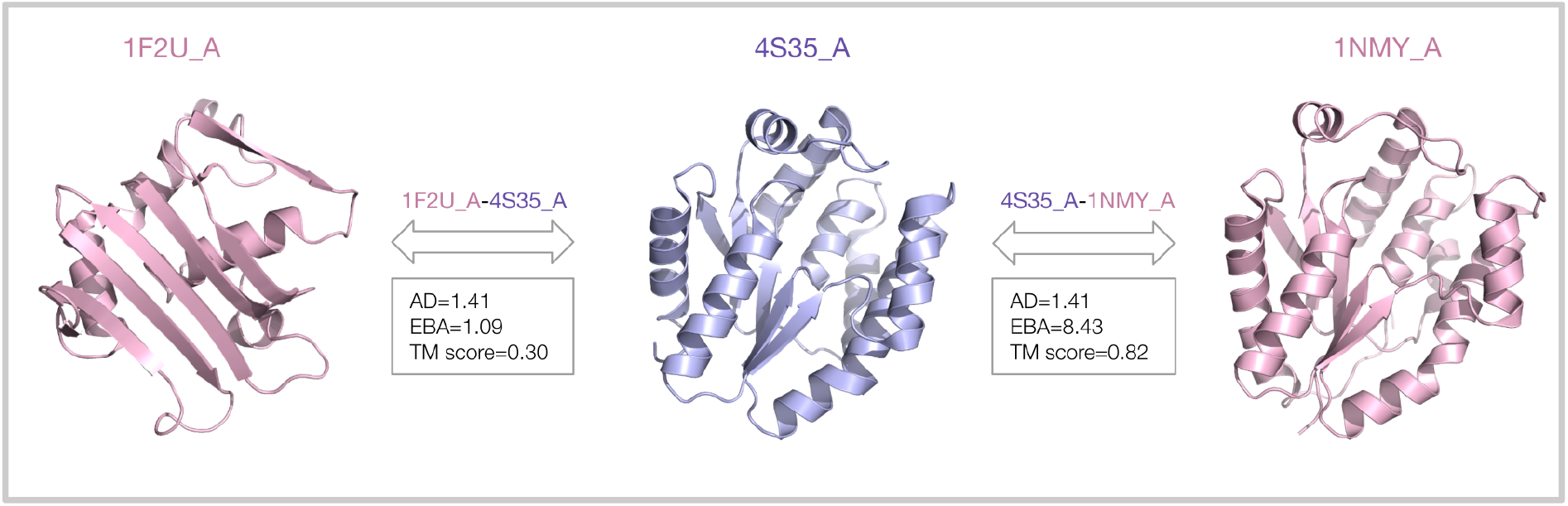
Comparison of three proteins - Thymidylate kinase from Aquifex Aeolicus VF5, human thymidylate kinase and Rad50 ATPase from Pyrococcus furiosus (PDB ids: 4S35, 1NMY and 1FU2 chain A) - using sequence and structure based scores. While 4S35 and 1NMY share a very similar fold (TM score=0.82), the 1F2U structure is different (TM score=0.30). However, the average representation of the protein sequence in the center (PDB id: 4S35) has approximately the same distance in the embedding space (AD=1.41) to both other proteins. In this example, the average distance (AD) is not able to distinguish the two cases, while our embedding-based alignment (EBA) assigns a much higher score to the pair of sequences with a similar fold. Notice that the proteins in this example share very little sequence identity: 15% for 1F2U-4S35 and 29% for 4S35-1NMY. Both the AD and EBA scores in this example were computed using the ProtT5 language model and the Euclidean distance metric.

In our work, we introduce an embedding-based alignment (EBA) approach that, given two sequences, leverages the distance of all possible pairs of residues to generate a “similarity matrix” that is then used as a score matrix in a classical dynamic programming alignment approach. The idea behind this approach is similar to [6], however the absence of any training and parameter optimization makes our method robust to generalization and easy to interpret. Furthermore, the method is not bound to a specific language model, therefore any pLM can be utilized, leaving a choice based on the requirements of specific scientific applications. The score obtained with this alignment is able to capture structural similarity even in the sequence-similarity twilight zone [16], outperforming other pLM methods and classic sequence based approaches in the detection of distant homologies. Such an approach allows the generation of reliable protein sequence and structure alignments at low sequence similarity, opening the door to the annotation and interpretation of protein sequences without clear homologs of known structure and function.

## 2 Methods

The methods described in this section rely on the assumption that residues with similar characteristics and context will have similar embeddings, therefore they will be close in the embedding space. Benchmarks have been performed with 2 pretrained state of the art pLMs: ProtT5-XL-UniRef50 (ProtT5) [4] and esm1b_t33_650M_UR50S (ESM-b1) [15]. Both models are based on the transformer architecture [22] and trained on UniRef50 [21] in a self-supervised fashion to predict masked amino acids. The residue-representations generated with these models are vectors belonging to spaces with, respectively, 1024 (ProtT5) and 1280 (ESM-b1) dimensions. It has been shown that, based on their position in the embedding space, amino-acids can be clustered according to biochemical and biophysical properties [4] [15].

### 2.1 Average distance-AD

Averaging per-residue embeddings is a simple and widely used approach to derive a fixed size representation for sequences of variable length [8], [7], [18]. Once the sequences are projected in this fixed size space, it is possible to compute the distance between them; we refer to this approach as the average distance (AD) method. Any distance metric can be used for this purpose and in this work, driven by preliminary analysis, we use Euclidean distances. AD is computationally efficient and captures meaningful relationships between proteins [8] [3] [18].

### 2.2 Embedding-based alignment-EBA

EBA aims to fully utilize the information encoded in the per-residue embeddings provided by pre-trained language models. Two protein sequences are compared by constructing a similarity matrix, which is used as score matrix to build an explicit alignment. The alignment score is finally used to define protein similarity. Given two sequences *A* and *B*, with lengths *n* and *m* respectively, the per-residue embedding similarity matrix *SM_n×m_* is built by computing the similarity score *SM_i,j_* for each possible pair of residues:

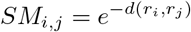

with *d*(): the desired distance metric, *r_i_*: embedding of residue *i* ∈ *A, r_j_*: embedding of the residue *j* ∈ *B*. All the analysis in this work were performed using Euclidean distance as *d*().

#### Signal enhancement

We will discuss in section 3.1 that a signal enhancement is beneficial for our method. The signal in the similarity matrix is enhanced by comparing the similarity of each pair of residues with the scores of all pairs involving the amino-acids of the interested couple. Given a pair of residues (i,j) with a similarity score *SM_i,j_*, we compute the Z-score with respect to both the elements in the same row (*SM*_*i*,*_) and column (*SM*_*,*j*_). We finally convert each element of the similarity matrix to the average of the computed Z-scores. In formalism:

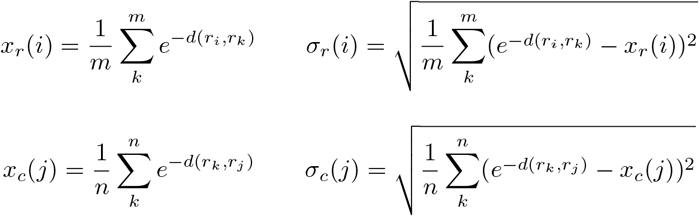

with *x*_*r/c*_(*i*/*j*) and *σ*_*r/c*_(*i/j*) being the average and standard deviation computed for the row/column i/j.

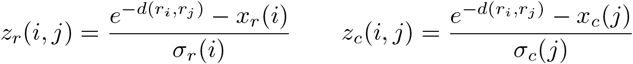

Each element of the enhanced similarity matrix is then computed as:

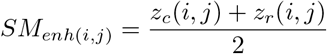

#### Dynamic alignment

A global gapped alignment with dynamic time warping [1] is performed using *SM_enh_* as score matrix and gap penalties set to 0. Given the resulting alignment score *s_align_*, the *EBA* similarity score is defined as:

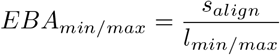

where *l_min/max_* is the length of the shorter/longer sequence involved in the comparison. The alignment score *s_align_* is symmetric with respect to the sequences *s_align_*(*A, B*) = *s_align_*(*B, A*). The symmetry is broken after normalization according to the length of one of the two, similarly to the normalization adopted for the computation of TM score for structure comparison [24].

### 2.3 Structural similarity analysis

We benchmarked AD and EBA in capturing structural similarities in the absence of clear sequence similarity. For that, we gathered protein pairs of known structure with low sequence identity using PISCES [23] (default parameters, with the exception of: ‘Maximum pairwise percent sequence identity’: 30% and ‘Minimum chain length’: 75). The resulting 19’599 pairs exhibit detectable homology (HHsearch [20] evalue threshold 10^-4^) but are only remotely related (sequence identity < 30%). Performances of EBA and AD were measured as Spearman correlation coefficients between the predicted similarity/distance and structural similarity, expressed as the TM score [25].

### 2.4 CATH annotation transfer analysis

We also benchmarked EBA against other pLM-based methods in transferring CATH domain annotations. In [7] annotations from a lookup set of 66K sequences were transferred to a test set of 219 CATH domains; in this work we reproduced their analysis using the same sets. The sequence identity between the test set and the lookup set is low and each sequence in the test set has at least one sequence with an identical label in the lookup set. Given a domain in the test set, the annotation of the domain with the higher EBA score across those in the lookup set is transferred. We carried out this analysis for each of the four CATH categories using EBA. We computed the accuracy of the annotation transfer as:

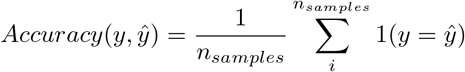

Results were compared to the scores reported in [7] for AD, ProtTucker [7] and HMMER [14].

## 3 Results

### 3.1 EBA captures structural similarity in the twilight zone

We benchmarked *EBA* against the following methods:

– *EBA* without signal enhancement (*EBA_plain_*) (section 2.2)
– AD (section 2.1)
– ProtTucker [7]
– TM-vec [6]
– Sequence identity provided by PISCES (SIP)[23]

We compared the predicted similarities/distances to the TM-scores computed with TM-align [25]. The asymmetric nature of TM-score allows to perform the analysis using both the score normalized by the length of the longer or the shorter sequence, *TM_min_* and *TM_max_* respectively. The choice of the score depends on the type of similarity one would like to investigate. The scores reported in table 1 shows the Spearman correlation computed using both *TM_min_* and *TM_max_*, normalizing the EBA score consistently with the TM-score normalization. Our results indicate that EBA outperforms all other approaches, independently of the underlying language model. Table 1 shows that *EBA* performs much better also than the not enhanced version: *EBA_plain_*. The strong impact of the signal enhancement suggests that raw distances, and thus the similarity scores, need to be contextualized as they are not necessarily comparable among different pairs of sequences. This is done by substituting the raw similarity of each residue pair with its pseudo Z-score, as described in section 2.2. This approach extracts the signal by assigning high values to residue pairs with high similarity with respect to other pairs involving the same residues.

**Table 1:** Spearman correlations between the similarity/distance predictions of the listed methods and the *TM* scores. When possible, we showcase the methods performances for both ProtT5 and ESM-b1. The *EBA* scores are normalized according to the TM scores, we therefore compare *EBA_min_* with *TM_min_* and *EBA_max_* with *TM_max_*. Since the other methods provide only one score, the same prediction is compared to both *TM_min_* and *TM_max_*. Notice that the expected correlation for similarity scores is positive, while for distances is negative.

| | $TM_{min}$ | | $TM_{max}$ | |
| --- | --- | --- | --- | --- |
|  | ESM-b1 | ProtT5 | ESM-b1 | ProtT5 |
| *EBA* | **0.87** | **0.90** | **0.80** | **0.84** |
| *EBA<sub>plain</sub>* | 0.64 | 0.72 | 0.52 | 0.54 |
| AD | -0.46 | -0.46 | -0.39 | -0.39 |
| ProtTucker |  | -0.46 |  | -0.38 |
| TM-vec |  | 0.78 |  | 0.70 |
| SIP | 0.21 |  | 0.26 |  |

### 3.2 Length normalization

The estimation of similarity between 2 proteins is affected by their difference in length. Whenever this difference is large, the choice of the normalization becomes an important factor. An example is shown in figure 2, where we consider a pair of sequences with the same length (pair 1) and a second pair in which one sequence is approximately double the size of the other (pair 2). In the first example, since the sequences have the same length, the normalization does not matter: *TM_min_* = *TM_max_* and *EBA_min_* = *EBA_max_*. In the second pair on the other hand, the shorter protein is completely contained in the longer one. In this case, *EBA_min_* and *EBA_max_* offer two different perspectives. The normalization according to the shorter sequence results in a large score (*EBA_max_* = 9.54), reflecting the fact that the shorter sequence successfully aligned though its whole length. However, the longer sequence is only partially aligned, therefore the normalization according to its length results in a lower score (*EBA_min_* = 4.41). Another way to tackle such a comparison would be the generation of a local alignment.

**Fig. 2:**
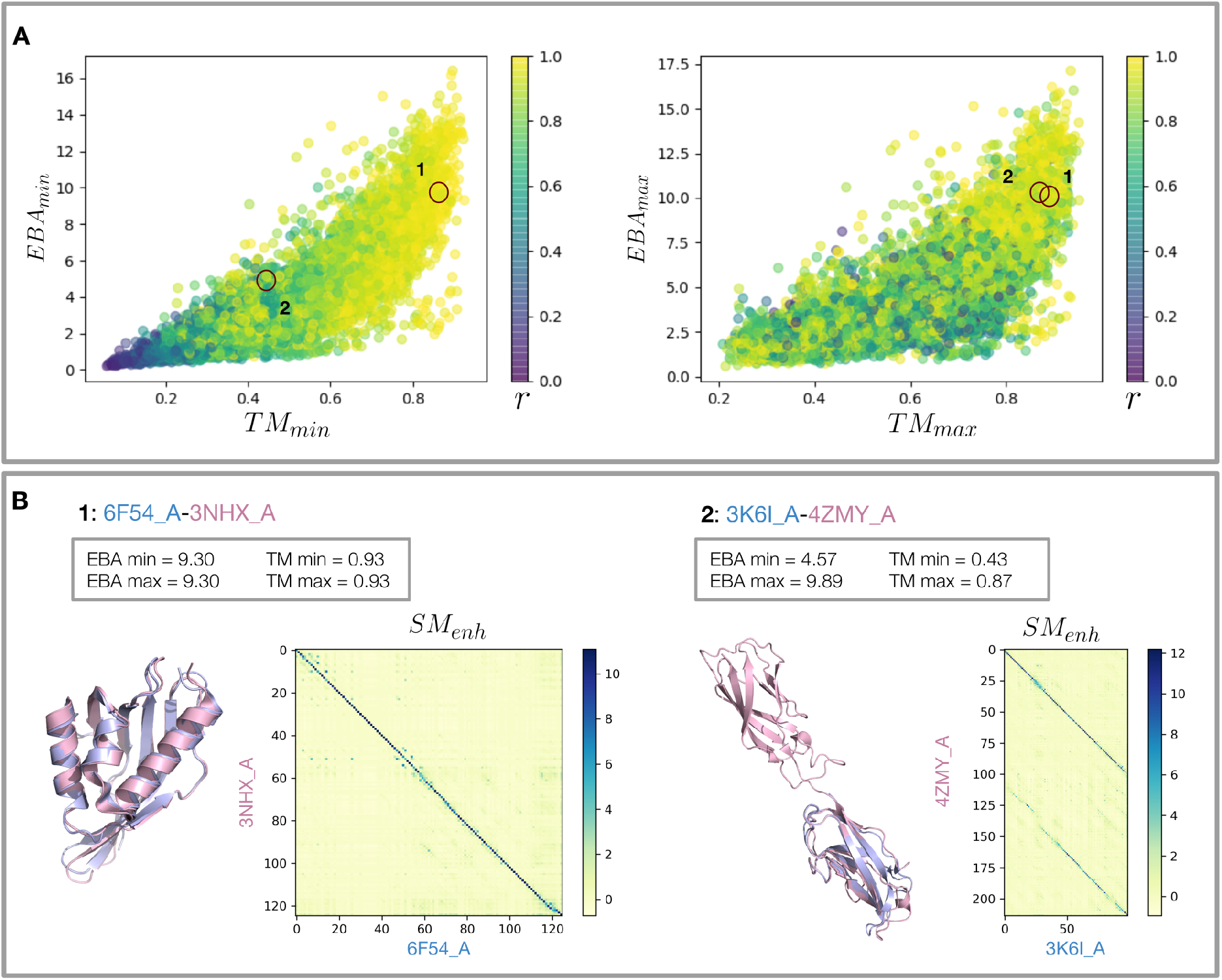
Panel A shows the correlation between *EBA_min/max_* and *TM_min/max_* for the analysis performed using ProtT5. A color gradient shows the length ratio of the sequence pairs: *r* = *l_min_*/*l_max_*. Where *l_min_* is the length of the shorter sequence and *l_max_* the length of the longer one. Panel B shows two pairs of sequences with very different *r*. Pair one is not affected by the length normalization, while pair 2 score changes drastically between *EBA_min_* and *EBA_max_*.

### 3.3 EBA vs AD

[18] already stressed the limitation of AD when comparing proteins with multiple domains. Here we illustrate another issue of AD: inherently losing order information that leads to the comparison of residues without any evolutionary relationship. In figure 3 we consider the example already displayed in figure 1. The scores for EBA and AD reported below are computed using ProtT5. While these two pairs of sequences exhibit similar AD (AD=1.41 for both), EBA favorably scores pair 1 (*EBA_min_* = 8.24) but not pair 2 (*EBA_min_* = 1.20). EBA thus better reflects structural similarity as measured by the TM score of the underlying structures (*TM_min_*: 0.82 and 0.25). The analysis of the pairwise residue-embedding distances (i.e., Euclidean distance in the 1024-dimensional embedding space) of the two sequence pairs helps us understand why EBA succeeds in distinguishing these two examples while AD fails. In the matrix representing pair 1 (figure 3 panel A), a diagonal is clearly visible. This diagonal not only shows how these 2 sequences have multiple residues close in embedding space, but also that they are ordered and thus alignable. In pair 2 (figure 3 panel B), on the other hand, despite having a matrix with overall similar numbers with respect to the first one (as suggested by the fact that the 2 pairs have similar AD) we do not observe any patterns of ordered residues close in embedding space. In figure 3 we also display the enhanced similarity matrix (*SM_enh_*) which strengthens the signal at the diagonal signal.

**Fig. 3:**
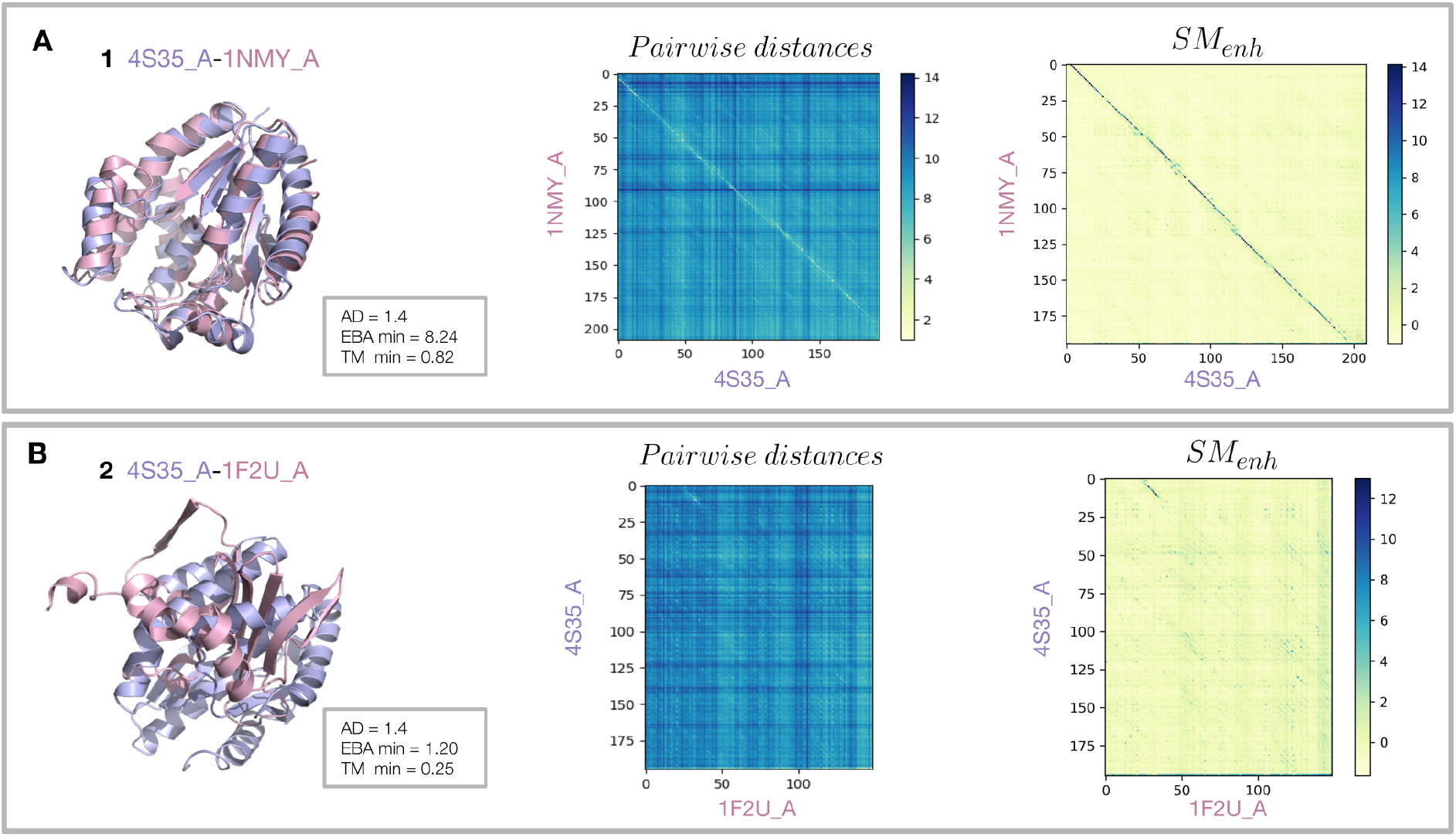
These are two pairs of sequences with very similar AD, but completely different TM scores. As we can see these are well classified from EBA. The distance matrices show the pairwise residue embedding distances of the two pairs. We can observe that in pair 1, a diagonal emerges from the plot. This shows how multiple ordered residues are close in the embedding space. While in pair 2, we do not observe any ordered signal. We also display the similarity matrix *SM_enh_* built for these two examples. We can see how the signal enhancement strengthens the signal for pair 1.

### 3.4 EBA successfully transfers CATH annotations

Table 2 shows a comparison between the best results reported in [7] against EBA. In this task, AD is outperformed by both EBA and ProtTucker. An expected result since the network underlying ProtTucker was trained for this specific purpose. EBA and ProtTucker offer similar performances. When using ProtT5, EBA is more accurate in transferring labels at the “Topology” and “Homology” levels, while ProtTuker performs better on the “Class” level. Although this test was performed on a small test set, our results indicate that EBA, which does not rely on training for a specific task nor on any parameter optimization, is able to successfully transfer CATH family annotations better than classic sequence profile based tools. As normalization for this analysis, we used the length of the longer sequence in each comparison, therefore *EBA_min_*. With this normalization, selecting the higher similarity scores ensures to value both similarity and sequence coverage in the comparison.

**Table 2:** Accuracy computed for the CATH annotation transfer analysis as in [7]. The reported *EBA* and *EBA_plain_* values are normalized according to the length of the longer sequence in each comparison: *EBA_min_*.

|  | ProtT5 |  |  |  | ESM-b1 |  |  | HMMER |
| --- | --- | --- | --- | --- | --- | --- | --- | --- |
|  | *EBA* | *EBA<sub>plain</sub>* | ProtTucker | AD | *EBA* | *EBA<sub>plain</sub>* | AD |  |
| C | 87 | 82 | 88 | 84 | **89** | 75 | 79 | 70 |
| A | 77 | 68 | 77 | 67 | **78** | 59 | 61 | 60 |
| T | **74** | 63 | 68 | 57 | 70 | 54 | 50 | 59 |
| H | **85** | 74 | 79 | 64 | 77 | 61 | 57 | 77 |
| average | **81** | 72 | 78 | 68 | 78 | 62 | 62 | 67 |

## 4 Conclusions

In this work, we showcase the potential of combining pLM representations and classical alignment methods for establishing distant homology relationships. Our embedding-based alignment (*EBA*) is able to identify structural similarities between proteins in the twilight zone, where pairwise sequence identity goes far below 30%. Our results indicate that, in such applications, pLM-based score matrices are a more robust option when compared to classic alternatives. This may be due to the ability of pLMs to capture not only residue biochemical characteristics, but also their context in the full proteins. Despite the absence of additional training or parameter optimization, *EBA* outperforms other state of the art pLM-based methods and classical sequenced-based approaches. The absence of any sort of re-training and optimization not only makes the approach extremely generalizable, but also allows to leverage different pLMs, which makes it adaptable to the extremely fast evolving field. However, *EBA* computation times are higher than the fast average-based methods. The average computation time for a pair-comparison is of the order 10^-2^ seconds on a CPU. While still reasonably fast for the alignment or comparison of small sets of sequences, this may become an issue in large-scale analyses. One way to overcome this is to carry out a pre-filtering step by first identifying putative close sequences using AD and then a higher-resolution alignment with *EBA*. This optimization approach was already proposed by similar works, such as: [6] and [10].

In our work we generated a global alignment, however the same score matrix (*SM_enh_*) can be employed for both a local alignment and multi-sequence alignment (MSA). A local alignment would help in the comparison of sequences of different lengths, as mentioned in 2, and would unlock other interesting possibilities, such as the identification of domain or circular permutations and the detection of repetitive patterns. On the other hand, an MSA approach would allow for the construction of MSAs involving highly divergent and dissimilar sequences, providing, for example, better inputs for deep learning methods that rely on MSAs, such as AlphaFold [9]. Recently, a pLM-based MSA method was proposed by [12]. Here, the authors generate MSAs by clustering and ordering amino acid contextual embeddings.

The rising popularity of embedding based alignment methods ([6], [12], [10]) highlights their potential. While writing this manuscript, multiple works on the topic became available. Particularly similar to ours is pLM-BLAST [10], where cosine similarities between per-residue embeddings are used as a proxy for measuring similarities of individual residue pairs. However, the biggest difference with *EBA* is probably the lack of signal enhancement. Therefore, pLM-BLAST is very similar to *EBA*_*plain*_, which is outperformed by *EBA*. This suggests that signal enhancing is an important factor when developing pLM-based protein alignment approaches.

We believe that the rapid development of such methods will soon further revolutionize protein bioinformatics; opening new doors into the modeling and annotation of proteins, beyond the detection horizon of current state-of-art tools.

## Data and code availability

The code to reproduce the analysis described in this paper is available at: https://git.scicore.unibas.ch/schwede/EBA. The repository also contains detailed instruction on how to generate the enhanced similarity matrix for a pair of sequences and score them with the EBA method.

## Acknowledgements

We would like to thank the SWISS-MODEL development team for insightful discussions, technical support and text revisions, and sciCORE at the University of Basel for providing computational resources and system administration support.

